# Anti-SARS-CoV-2 human antibodies retaining neutralizing activity against SARS-CoV-2 B.1.1.529 (omicron)

**DOI:** 10.1101/2021.12.21.473706

**Authors:** Maya Imbrechts, Winnie Kerstens, Madina Rasulova, Thomas Vercruysse, Wim Maes, Louanne Ampofo, Karen Ven, Jeroen Lammertyn, Karen Vanhoorelbeke, Nico Callewaert, Johan Neyts, Kai Dallmeier, Paul Declerck, Hendrik Jan Thibaut, Nick Geukens

## Abstract

SARS-CoV-2 B.1.1.529, designated omicron, was recently identified as a new variant of concern by WHO and is rapidly replacing SARS-CoV-2 delta as the most dominant variant in many countries. Unfortunately, because of the high number of mutations present in the spike of SARS-CoV-2 omicron, most monoclonal antibodies (mAbs) currently approved for treatment of COVID-19 lose their *in vitro* neutralizing activity against this variant. We recently described a panel of human anti-SARS-CoV-2 mAbs that potently neutralize SARS-CoV-2 Wuhan, D614G and variants alpha, beta, gamma and delta. In this work, we evaluated our mAb panel for potential *in vitro* activity against SARS-CoV-2 delta and omicron. Three mAbs from our panel retain neutralizing activity against both delta and omicron, with mAb 3B8 still resulting in complete neutralization at a concentration as low as 0.02 μg/ml for both variants. Overall, our data indicate that mAb 3B8 may have the potential to become a game-changer in the fight against the continuously evolving SARS-CoV-2.

## Introduction

The latest SARS-CoV-2 variant of concern (VoC) B.1.1.529, designated omicron, was first reported to WHO from South Africa at the end of November and is rapidly spreading across many countries, thereby replacing the already highly transmissible SARS-CoV-2 delta (B.1.617.2) VoC. SARS-CoV-2 omicron has a large number of mutations compared to its pandemic founder, many of which are predicted to result in resistance towards the currently available neutralizing (therapeutic) antibodies, including those induced upon vaccination^1,2^. Teams all over the world are racing to test the activity of their antibodies against SARS-CoV-2 omicron, with the majority of them showing strongly reduced or no activity. To the best of our knowledge, only GSK has reported that their therapeutic monoclonal antibody (mAb) Sotrovimab retains activity against the omicron variant (IC_50_ of 0.27 μg/ml)^3–5^, whereas mAbs REGN10933 and REGN10987 from Regeneron, and Ly-CoV016 and Ly-CoV555 from Abcellera/Eli Lilly, all failed to neutralize omicron at the highest concentration tested (10 μg/ml) in a pseudovirus neutralization assay^4^. In addition, also the lead mAb from Adagio Therapeutics, which was specifically designed to be highly potent and broadly neutralizing, showed a more than 300-fold reduction in neutralizing capacity against omicron when tested in both an authentic and pseudovirus assay^6^.

We recently described a panel of eight human anti-SARS-CoV-2 RBD-specific mAbs, derived from a convalescent COVID-19 patient, potently neutralizing SARS-CoV-2 D614G (manuscript under review)^7^. Six of these mAbs retained activity against SARS-CoV-2 Wuhan, alpha, beta, gamma and delta infection *in vitro*. In hamsters, two mAbs (3E6 and 3B8) potently cured infection with SARS-CoV-2 Wuhan, beta and delta when administered therapeutically at 5 mg/kg. 3B8 was remarkably potent and still reduced lung SARS-CoV-2 delta viral titers at a dose as low as 0.2 mg/kg^7^. Building on these previous data, we now tested our mAb panel against SARS-CoV-2 omicron *in vitro*.

## Results

We evaluated the neutralizing capacity of our eight anti-SARS-CoV-2 human mAbs against SARS-CoV-2 delta and omicron, using a pseudovirus assay with vesicular stomatitis virus (VSV) expressing the SARS-CoV-2 spike protein from the delta or omicron variant. As shown in Figure 1A, six out of eight mAbs neutralized SARS-CoV-2 delta, with mAb 3B8 still resulting in complete neutralization at a concentration as low as 0.02 μg/ml (Table 1), similar to our previously reported data using a plaque reduction neutralization assay^7^. In addition, three of the eight mAbs also retained neutralizing activity towards SARS-CoV-2 omicron (Figure 1B and Table 1). mAbs 2B2 and 1A10 showed 50% neutralization of omicron (IC_50_) at a concentration of 1.1 and 1.7 μg/ml respectively. Highly interesting, mAb 3B8 still completely neutralized SARS-CoV-2 omicron even at the lowest concentration tested (0.02 μg/ml).

**Figure 1:**
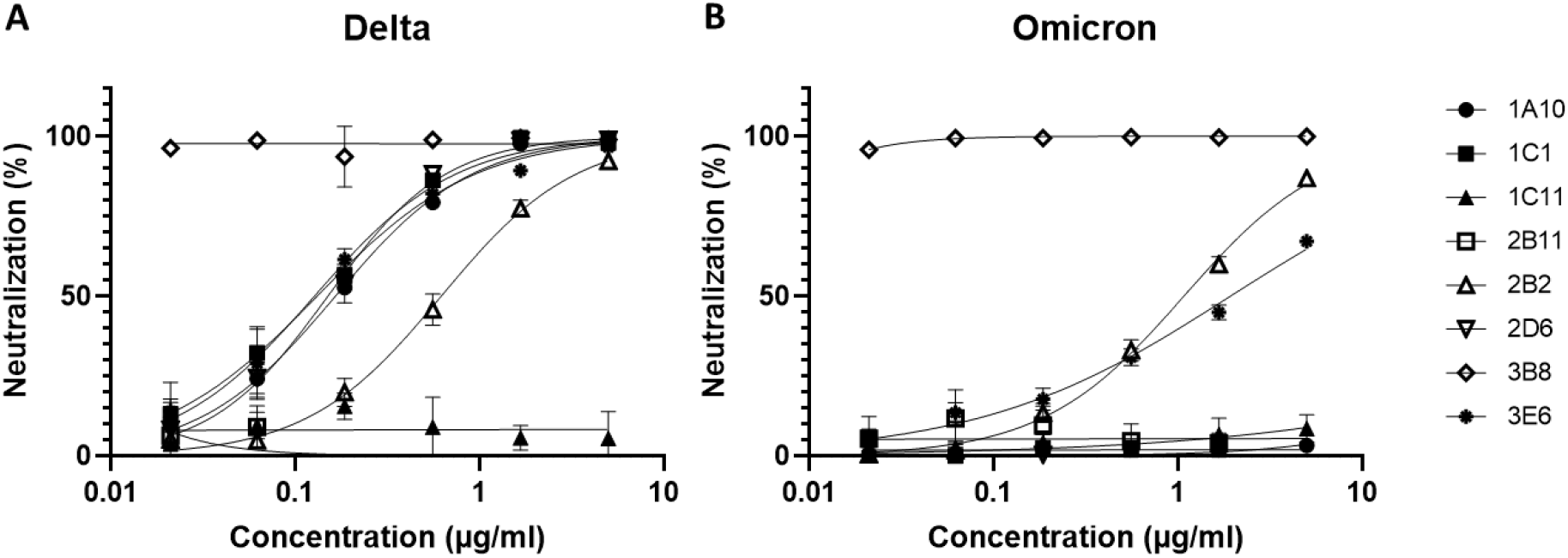
Neutralizing activity of mAbs against SARS-CoV-2 delta and omicron VoC. Percentage neutralizing activity, measured via a SARS-CoV-2 delta (A) or omicron (B) VSV pseudotyped assay, at different concentrations of the respective mAbs tested. Symbols show mean and standard deviation of 3 replicates, measured in one experiment.

**Table 1:** Overview of *in vitro* neutralizing activity (IC_50_) towards SARS-CoV-2 delta and omicron.

| | $IC_{50}$ ($\mu\text{g/ml}$) | |
| --- | --- | --- |
|  | Delta | Omicron |
| **1A10** | 0.18 | ND |
| **1C1** | 0.12 | ND |
| **1C11** | ND | ND |
| **2B11** | ND | ND |
| **2B2** | 0.54 | 1.11 |
| **2D6** | 0.14 | ND |
| **3B8** | <0.02 | <0.02 |
| **3E6** | 0.13 | 1.69 |

Neutralizing activity ( $IC_{50}$ ; in $\mu\text{g/ml}$ ) was determined via a VSV pseudotyped assay for SARS-CoV-2 delta and omicron. ND = no detectable neutralizing activity at maximum tested concentration of 5 $\mu\text{g/ml}$ .

Experiments to fully explore the activity range of 3B8 against SARS-CoV-2 delta and omicron are currently ongoing.

## Discussion

In this work, we report on the *in vitro* neutralizing activity against pseudotyped SARS-CoV-2 delta and omicron of eight mAbs recently described by our group. We identified three mAbs that retain neutralizing activity against both delta and omicron, with mAb 3B8 showing a very high potency with complete neutralization at a concentration of 0.02 μg/ml for both variants. Although data are from only one experiment with triplicate measurements, the results reported here for neutralization of SARS-CoV-2 delta are in line with those previously described using another assay format (VSV pseudotyped assay versus plaque reduction assay respectively), thereby indicating a high reliability of the current data. Additional experiments *in vitro* as well as *in vivo* need to be performed to confirm the mAb’s neutralizing activity against SARS-CoV-2 omicron. Of note, as our mAb panel was isolated from a convalescent COVID-19 patient who got infected with SARS-CoV-2 in March 2020 (and therefore probably with SARS-CoV-2 Wuhan), our data also show that at least some of the neutralizing antibodies elicited upon infection (or vaccination) with the Wuhan strain potently cross-react with SARS-CoV-2 omicron. Considering the broad cross-reactivity and high potency of mAb 3B8 reported previously^7^, together with its strong neutralizing capacity towards SARS-CoV-2 omicron *in vitro* described here, in combination with the fact that multiple currently available therapeutic mAbs lose their activity against omicron, we believe 3B8 might be a game-changer in the fight against the continuously evolving SARS-CoV-2.

## Material and methods

### Production of S-pseudotyped virus and serum neutralization test

VSV S-pseudotypes were generated as described previously^8^. Briefly, HEK-293T cells were transfected with the respective S protein expression plasmids for the delta variant (Cat. No. plv-spike-v8, Invivogen) and the omicron variant (A67V, Δ69-70, T95I, G142D, Δ143-145, Δ211, L212I, ins214EPE, G339D, S371L, S373P, S375F, K417N, N440K, G446S, S477N, T478K, E484A, Q493R, G496S, Q498R, N501Y, Y505H, T547K, D614G, H655Y, N679K, P681H, N764K, D796Y, N856K, Q954H, N969K, L981F as based on the information available on the ECDC website on November 29^th^, 2021) and one day later infected (MOI = 2) with GFP-encoding VSVΔG backbone virus (purchased from Kerafast). Two hours later, the medium was replaced by medium containing anti-VSV-G antibody (I1-hybridoma, ATCC CRL-2700) to neutralize residual VSV-G input. After 24 h incubation at 32 °C, the supernatants were harvested. To quantify neutralizing mAbs, serial dilutions of serum samples were incubated for 1 hour at 37 °C with an equal volume of S pseudotyped VSV particles and inoculated on Vero E6 cells. In parallel to the mAbs, human convalescent serum (WHO standard NIBSC 20/130; stock concentration 1300 IU/ml; start dilution 1/400) was included as benchmark.

The percentage of GFP expressing cells was quantified on a Cell Insight CX5/7 High Content Screening platform (Thermo Fischer Scientific) with Thermo Fisher Scientific HCS Studio (v.6.6.0) software. Neutralization IC_50_ values were determined by normalizing the serum neutralization dilution curve to a virus (100%) and cell control (0%) and fitting in Graphpad Prism (GraphPad Software, Inc.) (inhibitor vs. response, variable slope, four parameters model with top and bottom constraints of 100% and 0% respectively).

## Acknowledgements

We would like to thank Jasmine Paulissen, Catherina Coun and Nathalie Thys (TPVC) for skilled and rapid generation of spike-pseudotyped virus and diligent serology assessment.

This work was funded by KU Leuven (project SARS-CoV-2 mAb-OF2 and the Covid-19-Fund KU Leuven/University Hospital Leuven), the COVID-19 call of Fund for Scientific Research Flanders (FWO) (grant G0G4820N), the European Union’s Horizon 2020 research and innovation program under Grant Agreement 101003627 (Swift COronavirus therapeutics REsponse project) and Bill and Melinda Gates Foundation under Grant Agreement INV-00636.

## References

1. World Health Organization. Classification of Omicron (B.1.1.529): SARS-CoV-2 Variant of Concern. 26/11/2021 (2021).

2. CDC. SARS-CoV-2 Variant Classifications and Definitions. https://www.cdc.gov/coronavirus/2019-ncov/variants/variant-info.html#Interest (2021).

3. GlaxoSmithKline. Preclinical studies demonstrate sotrovimab retains activity against the full combination of mutations in the spike protein of the Omicron SARS-CoV-2 variant. 7/12/2021 https://www.gsk.com/en-gb/media/press-releases/sotrovimab-retains-activity/ (2021).

4. Sheward, D. J. et al. Variable loss of antibody potency against SARS-CoV-2 B.1.1.529 (Omicron). bioRxiv Prepr. Serv. Biol. (2021).

5. Cathcart, A. L. et al. The dual function monoclonal antibodies VIR-7831 and VIR-7832 demonstrate potent in vitro and in vivo activity against SARS-CoV-2. bioRxiv 2021.03.09.434607 (2021) doi:10.1101/2021.03.09.434607.

6. Adagio Therapeutics. Adagio Therapeutics Reports Reduction in In Vitro Neutralizing Activity of ADG20 Against Omicron SARS-CoV-2 Variant. 14/12/2021 (2021).

7. Imbrechts, M. et al. Potent neutralizing anti-SARS-CoV-2 human antibodies cure infection with SARS-CoV-2 variants in hamster model. bioRxiv Prepr. Serv. Biol. (2021).

8. Sanchez-Felipe, L. et al. A single-dose live-attenuated YF17D-vectored SARS-CoV-2 vaccine candidate. Nature 590, 320–325 (2021).

